# X10 Expansion Microscopy Enables 25 nm Resolution on Conventional Microscopes

**DOI:** 10.1101/172130

**Authors:** Sven Truckenbrodt, Manuel Maidorn, Dagmar Crzan, Hanna Wildhagen, Selda Kabatas, Silvio O. Rizzoli

## Abstract

Expansion microscopy is a recently introduced imaging technique that achieves super-resolution through physically expanding the specimen by ~4x, after embedding into a swellable gel. The resolution attained is, correspondingly, approximately 4-fold better than the diffraction limit, or ~70 nm. This is a major improvement over conventional microscopy, but still lags behind modern STED or STORM setups, whose resolution can reach 20-30 nm. We addressed this issue here by introducing an improved gel recipe that enables an expansion factor of ~10x in each dimension, which corresponds to an expansion of the sample volume by more than 1000-fold. Our protocol, which we termed X10 microscopy, achieves a resolution of 25-30 nm on conventional epifluorescence microscopes. X10 provides multi-color images similar or even superior to those produced with more challenging methods, such as STED, STORM and iterative expansion microscopy (iExM), in both cell cultures and tissues.

---

The resolution of fluorescence microscopes has been limited by the diffraction barrier to approximately half of the wavelength of the imaging light (in practice, 200-350 nm). This barrier has been lifted by several microscopy concepts, for example by using patterned light beams to determine the coordinates from which fluorophores are permitted to emit, as in the stimulated emission depletion (STED) family,^1,2^ or by determining the positions of single fluorophores that emit randomly, as in photo-activated localization microscopy (PALM),^3^ stochastic optical reconstruction microscopy (STORM and dSTORM),^4,5^ or ground state depletion microscopy followed by individual molecule return (GSDIM).^6^ Although such technologies have been applied to biology for more than a decade, their general impact on biomedical research is still relatively limited. This is mainly due to the fact that accurate super-resolution is still available only for selected laboratories that are familiar with the different tools, are able to apply the appropriate analysis routines, and/or possess the often highly expensive machinery required.

The ideal super-resolution tool for the general biologist needs to be easy to implement, without specialized equipment, and without the need for complex imaging analysis. At the same time, such a technique would need to be highly reliable, and should be easy to apply to multiple color channels simultaneously. The expected resolution should be at least on the size scale of the labeling probes used. This would be ~20-30 nm for normal immunostaining experiments, since these rely on identifying the epitopes via primary antibodies that are later detected through secondary antibodies, each of which are ~10-15 nm in size. Expansion microscopy, a technique introduced by the Boyden laboratory,^7-10^ is an important step in this direction. Expansion microscopy entails that the sample of interest is first fixed, permeabilized and immunostained, and is then embedded in polyelectrolyte gels which expand strongly when dialyzed in water. To ensure that no disruption of the sample aspect ratio occurs, the sample is digested using proteases after embedding, but before the expansion step. The fluorophores, which are covalently bound to the gel, thus maintain their relative positions, although they are now positioned a few-fold farther away from each other than in the initial sample. The preparation can then be imaged in a conventional microscope. This renders expansion microscopy the simplest approach, to date, that is able to produce super-resolution images. However, the initial implementations of this approach were performed with gels that expanded, on average, about 4-fold, which resulted in lateral resolutions of ~70 nm, *i.e.* not as high as that of modern STED or STORM microscopes.^7^ The only solution proposed so far to this problem has been to use complex procedures consisting of multiple successive expansion steps (iterative expansion), which would require the embedding, expansion, re-embedding, and re-expansion of the sample.

We set out here to solve this problem, by generating a protocol that uses only one embedding and expansion step, but still obtains a resolution of the required value (20-30 nm), in multiple color channels, without any difficult techniques, tools or analysis routines. Our protocol expands the sample by 10-fold, and we therefore termed it X10 microscopy. It achieves a resolution of 25-30 nm on conventional epifluorescence microscopes, and does not even require confocal imaging for accurate nanoscale analyses. We compared X10 microscopy with state-of-the-art commercial implementations of both STED and STORM, and found it to be superior to both. Judging from the available literature, it is clear that self-build super-resolution microscopes, operated and optimized by specialists, could provide images that are superior to our X10 implementations on epifluorescence setups. In spite of this, the fact that X10 is the simplest and cheapest super-resolution technique currently available, with a resolution performance that is superior to what is available to the general biologist (*i.e.* the commercial implementations of these techniques), should render it the tool of choice for the implementation of super-resolution in the general biology laboratory.

## RESULTS AND DISCUSSION

To obtain a resolution of 20-30 nm within a one-step expansion procedure, we generated a protocol that enables the use of a superabsorbent hydrogel designed for excellent mechanical sturdiness^11^ for the expansion of biological samples. This gel uses *N,N*-dimethyl-acrylamide (DMAA) for generating polymer chains, which are crosslinked with sodium acrylate (SA) to produce a swellable gel matrix (Figure 1). The gelation free radical polymerization reaction is catalyzed by potassium persulfate (KPS) and *N,N,N’,N’*-tetramethyl-ethylene-diamine (TEMED; Figure 1A), and produces a gel that can expand >10x in each dimension when placed in distilled water (Figure 1B-C). Protein retention in the gel is achieved via the previously described anchoring approach,^8,10^ by employing Acryloyl-X. This uses NHS-ester chemistry to covalently attach to proteins, while a second reactive acrylamide group integrates into the polymerizing gel matrix.

**Figure 1.**
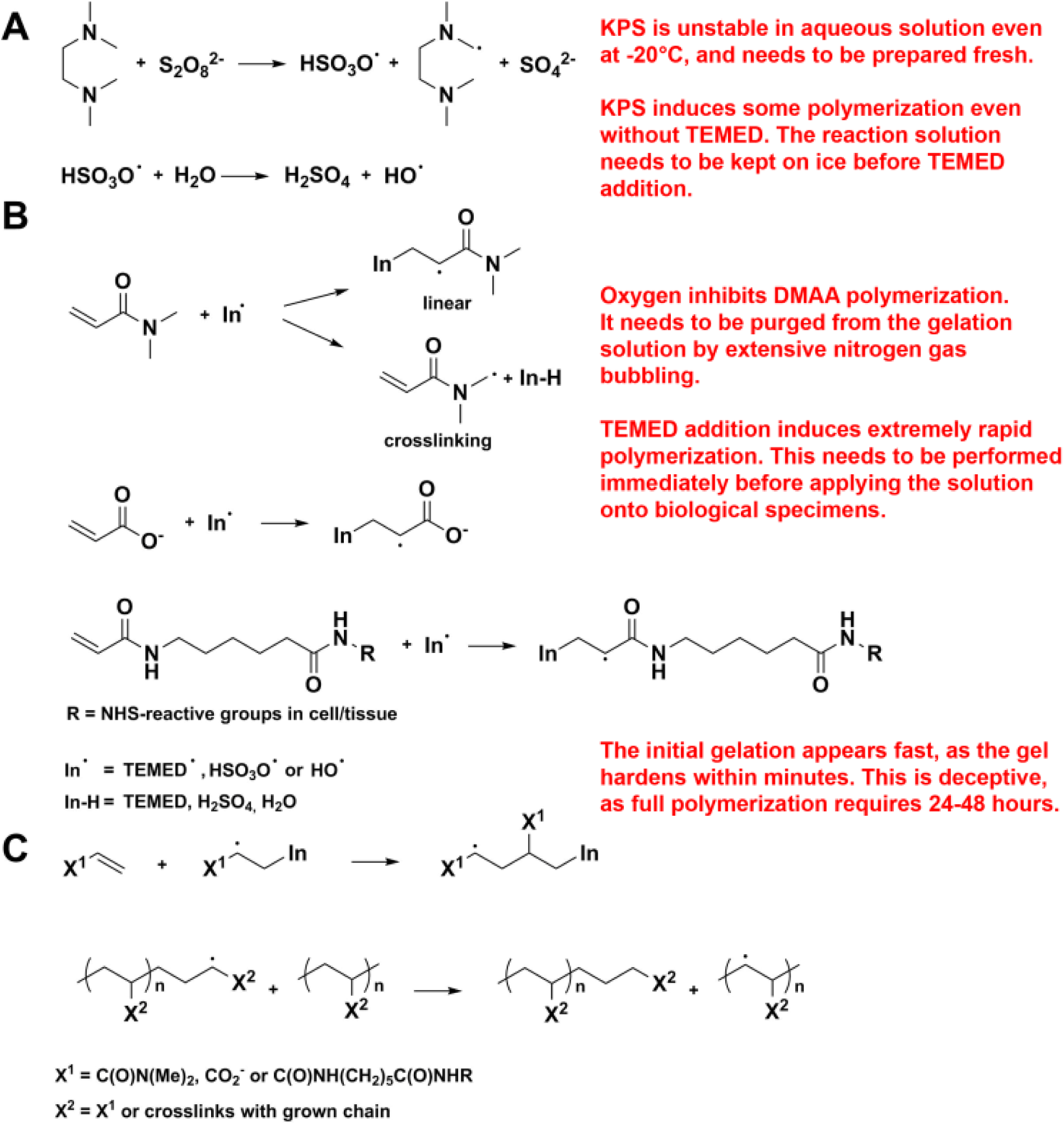
X10 gel polymerization reactions. (A) Primary TEMED, sulfate, and hydroxyl radicals are generated by redox initiation with KPS and TEMED.^32^ (B) Radical propagation occurs when the monomer (DMAA),^11,33^ ionic co-monomer (SA),^11^ and tissue-anchored acryloyl-monomer react with the primary radicals. Besides linear growth of the resulting polymer, DMAA also cross-links after proton abstraction at the methylene group.^11,33^ (C) The radical chain grows by reacting with monomers, and through radical transfer to monomers or other polymers to form a branched network. Critical steps are shown in red.

The different steps of the gel formation and protein retention reactions were initially difficult to optimize, and therefore required extensive testing and fine-tuning. Nevertheless, the final version of the protocol is trivially simple, and is highly reproducible. We present the critical steps in red in Figure 1, and we have included a complete protocol in the Supplementary Methods. Briefly, the main issues are the following. First, the reactions are extremely fast, and therefore low temperature and high speed of application are essential, unlike in the gels used for 4x expansion. Second, oxygen inhibits polymerization, and therefore needs to be carefully eliminated by bubbling with N_2_. This is a trivial procedure, which requires no specialized setup (other than a tube to conduct the N_2_ gas from a pressured gas container into the reaction mixture). Third, the gelation is initially rapid (it only takes minutes for the initial hardening), but does not continue with the same speed, and therefore care must be taken that the gel is allowed to polymerize for 24-48 hours.

Should these steps be followed as described here, the resulting gel formation is highly reproducible (Figure 2, and Figure S2). The maximum expansion factor we achieved with this approach was ~11.5x (Figure 2A,C; see also Supplementary Methods), which results in images with an apparent lateral resolution of ~25-30 nm (predicted from Abbe’s resolution limit; Figure 2A-C and Figure 3), in which substantially more details are revealed (Figure 2A-C, and Movie S1).

**Figure 2.**
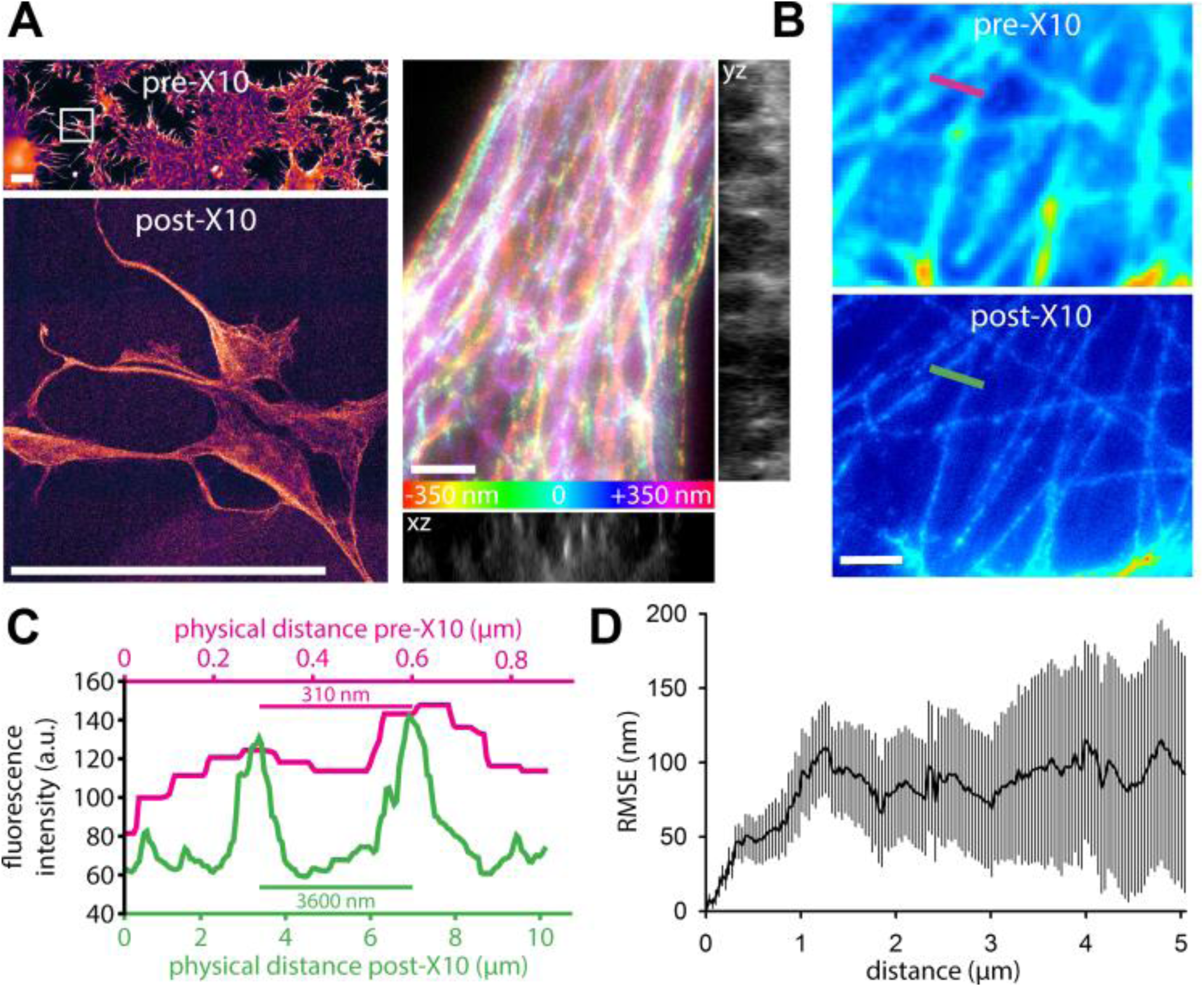
X10 achieves a resolution of 30 nm on conventional epifluorescence microscopes. (A) The X10 gel is swellable to >10x of its original size. The top panel on the left shows an overview image of COS7 cells stained for Tubulin, before expansion. The bottom panel shows the cells framed in the top panel (white rectangle), after expansion. The images are to scale, demonstrating an expansion factor of 11.4x in this example. Scale bars: 100 μm (both panels). The post-expansion image is dimmer, as the fluorophores are diluted ~1000-fold, and therefore requires a longer acquisition time and a higher camera gain. The right panels reveal the 3D organization of the Tubulin network in COS7 cells. The relative axial position of the fluorophores is visualized in a z-stack projection by colour-coding (see scale at the bottom). Orthogonal views are given next to the z-stack projection (yz view across the midline, xz view along the bottom). A movie through this z-stack, including a rocking projection, is available in Movie S1. Expansion factor: 11.4x. Scale bar: 1 μm. (B) Comparison between pre-expansion resolution of Tubulin imaging in COS7 cells (left panel) and post-expansion resolution in the same sample (right panel). Note that the images have not been processed to minimize distortions or to achieve a better correlation. Expansion factor: 11.5x. Scale bar: 1 μm. (C) An exemplary measurement for the X10 expansion factor. A line scan was drawn over corresponding regions before and after expansion, as indicated in panel B) by the coloured lines. (D) An analysis of the root mean square error (RMSE) of the distortions between aligned pre - and post-expansion images (see Supplementary Methods for details; n = 34 automated measurements from 4 independent experiments).

The resulting technique is fully compatible with the use of common affinity probes, such as antibodies (Figure 2), since X10 requires no specially designed labelling tools, similar to recent improvements to the 4x expansion.^8,12^ The distortions of the sample introduced by the gel during swelling are minimal (Figure 2D), and are virtually identical to those seen in 4x expansion microscopy.^7,8,12^ We would like to note, however, that the extensive digestion required for X10 is incompatible with expansion microscopy protocols that preserve fluorescent proteins.^10^ These protocols utilize a milder digestion that retains some fluorescent proteins. This milder digestion, however, does not allow X10 to retain the sample integrity at higher expansion factors (Figure S1). Therefore, fluorescent proteins will be visualized in X10 only by immunostaining them. However, this is not a major difficulty, as antibodies are currently available for all major fluorescent proteins. We would also like to note that X10 once more highlights the need for new probes for super-resolution imaging, as conventional antibodies usually do not result in a continuous staining of microtubules, but in a pearls-on-a-string pattern (as visible in Figure 2). This artefact, which is due to incomplete epitope coverage through conventional antibodies,^13,14^ can be observed also in many published works using other super-resolution techniques, such as STED^14,15-18^ and STORM^17,19-22^ (see also Figure S2). Alternatively, highly optimized tubulin labelling protocols should be used, to ensure optimal epitope coverage.

To verify the resolution of X10 experimentally, we relied on investigating peroxisomes, which are round organelles with dimensions of ~100-200 nm in neurons. We immunostained Pmp70, a protein of the peroxisome membrane (Figure 3), and we compared pre-expansion images with post-expansion images, as well as with STED and STORM images (Figure 3A; see Figure S3 for a more detailed comparison, and Movie S2 for a z-stack through several peroxisomes). To determine the nominal resolution of X10, we drew line scans through the membranes of the peroxisomes (post-expansion), fitted them to Gaussian curves, and determined their full width at half maximum values (FWHM; Figure 3B). The resolution determined in this fashion fits the theoretical prediction from Abbe’s resolution limit that we have stated above, being centered at 25.2 ± 0.2 nm (Figure 3C).

**Figure 3.**
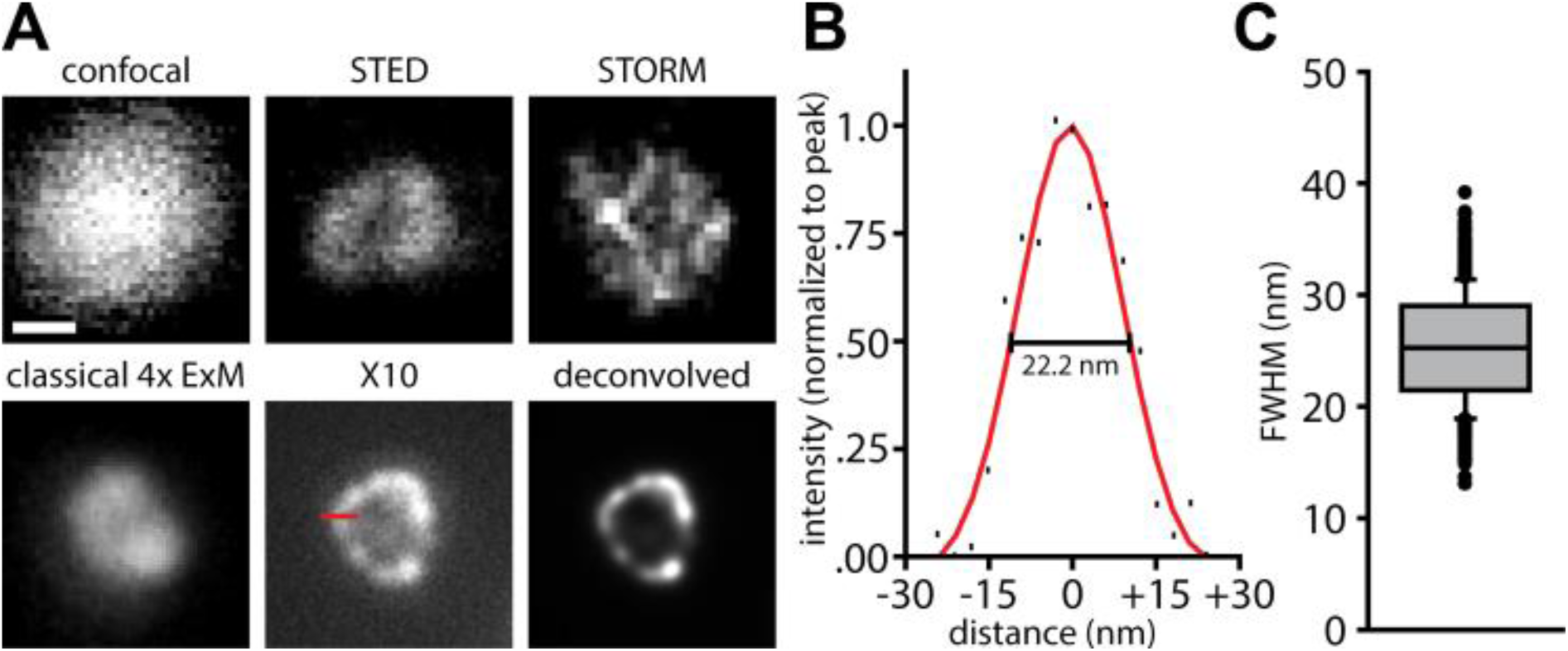
The resolution of X10 is ~25 nm. (A) Immunostainings for the peroxisome membrane protein Pmp70 in neurons are shown. The first 5 panels show individual peroxisomes imaged with a confocal microscope before expansion, with a STED microscope before expansion, with a STORM microscope before expansion, with an epifluorescence microscope after classical 4x expansion microscopy, and with an epifluorescence microscope after X10 (without and with deconvolution). Expansion factors: 3.8x for classical 4x expansion microscopy and 9.5x for X10. Scale bar: 100 nm (applies to all panels). The red line in the X10 panel indicates a line scan over the peroxisome membrane (60 nm in length). See Figure S3 for further examples. (B) The exemplary line scan from the X10 image in A) is shown with a best Gaussian fit curve, with an indicated measurement of resolution as full width at half maximum (FWHM). (C) A quantification of the average resolution, which is 25.2 ± 0.2 nm (n = 653 line scans across peroxisomes from 2 independent experiments).

We have also simulated peroxisomes stained for Pmp70, taking into account the size and random orientation of the primary/secondary antibody complexes (Figure S4; see Supplementary Methods for details), and found that the measured resolution value fits closely to the one predicted by the simulations (22.8 nm, on 10,000 simulated peroxisomes). This level of resolution is usually only achieved in highly specialized applications of STED and STORM microscopy,^23,24^ or in iterative expansion microscopy (iExM),^25^ and is bettered substantially only by a recently developed tool, MINFLUX microscopy.^26^ When investigating the same protein in state-of-the art commercial STED and STORM setups, the image quality these techniques achieved was, at best, comparable to that of X10 (Figure 3A; Figure S3). We used the modelling approach to determine if we could, theoretically, resolve the lumen of microtubules, but found that this is beyond the limits of X10, when implemented with epifluorescence microscopy (Figure S5), due to problems in the placement of antibodies across the expanded microtubule. Their large size effectively limits resolution,^25,27^ which implies that the ~25 nm resolution is possibly the maximum useful resolution that can be achieved in expansion microscopy when using conventional primary/secondary antibody stainings and epifluorescence microscopy.

The X10 procedure can be used to achieve multi-colour super-resolution imaging. We could easily resolve, for example, synaptic vesicle clusters in cultured hippocampal neurons, along with the presynaptic active zones and the postsynaptic densities (Figure 4A-C). This enabled us to measure the distance between the presynaptic active zone, identified by Bassoon, and the postsynaptic density, identified by Homer 1 (Figure 4D). We found this distance to be ~120-140 nm, very similar to what has been previously described for these proteins using STORM microscopy.^23,28^ The morphology of these structures is also very similar to what has been observed with advanced STORM microscopy.^29^ This type of information could not be obtained with either conventional microscopy or with classical 4x expansion microscopy (Figure S6).

**Figure 4.**
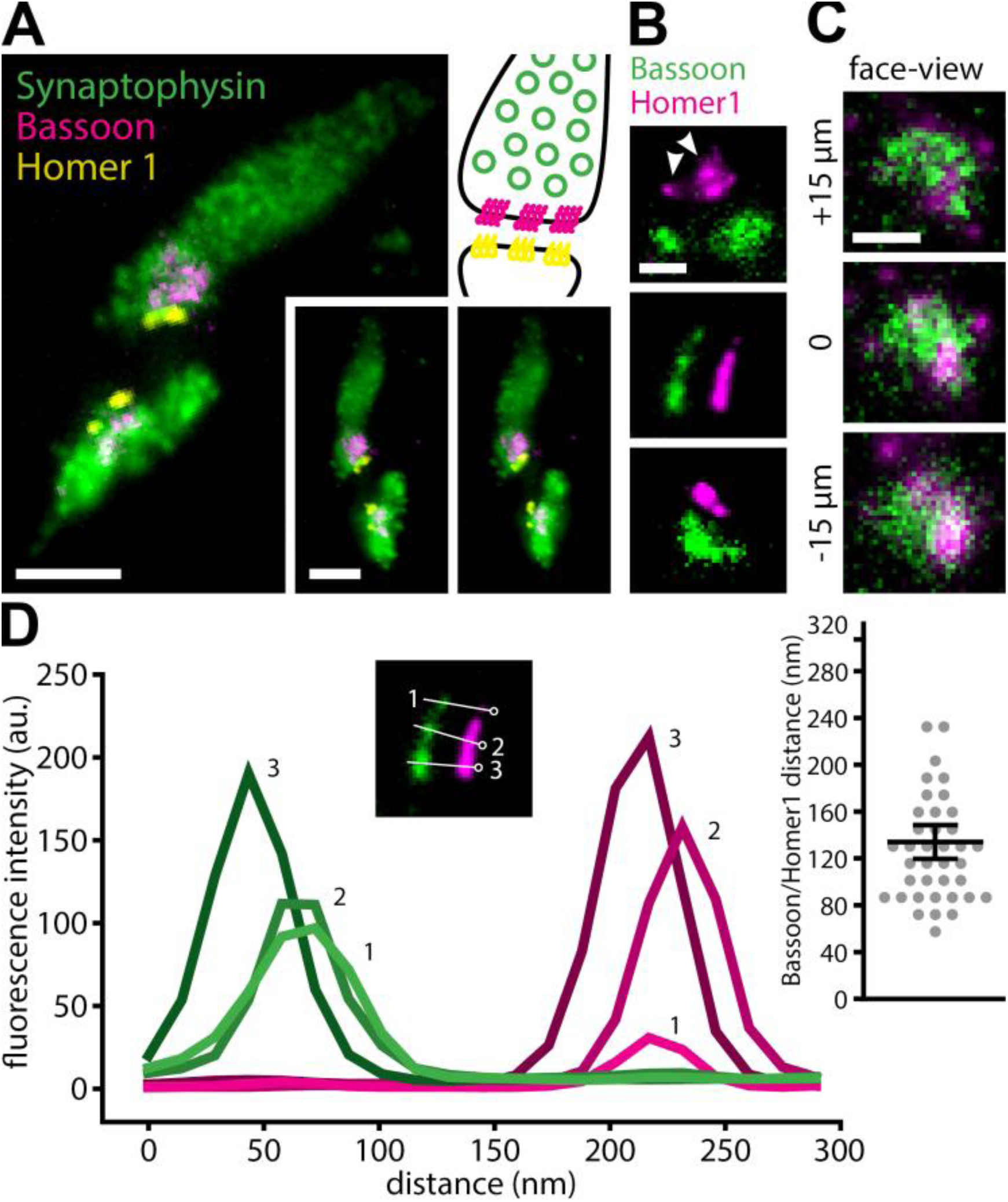
Multi-colour imaging with X10 reveals synaptic ultrastructure in cell culture. (A) 3-colour imaging resolves synaptic vesicle clusters (identified by Synaptophysin), along with presynaptic active zones (identified by Bassoon) and postsynaptic densities (identified by Homer 1). The panel at the top right gives a schematic overview of the organization of a synapse, for orientation (colours as in the fluorescence images). The 2 panels on the bottom right provide a stereo view of the synapses. Expansion factor: 11.0x. Scale bars: 500 nm (both). (B) Higher-magnification images show the alignment of presynaptic active zones and postsynaptic densities, as well as the distance between them, in side-view. Expansion factor: 11.0x. Scale bar: 200 nm. (C) A z-stack through an additional synapse, in face-view. Expansion factor: 11.0x. Scale bar: 200 nm. (D) Line scans through presynaptic active zones and through the corresponding postsynaptic densities reveal the distance between the two. The image inset shows three example line scans, and identifies them by number (green is Bassoon, magenta is Homer 1). The inset on the right quantifies the range of observed distances (n = 38 active zones and corresponding postsynaptic densities; mean ± SEM are 127 ± 7.1, indicated by the black lines).

Overall, such examples demonstrate that X10 microscopy can reproduce results that were previously obtained only with highly specialized imaging tools. In addition, the ease with which multiple colours can be investigated in X10 is an advantage over previous localization microscopy methods, which have been typically limited to two colour channels in practice. Importantly, X10 microscopy can also be applied to thin tissue slices, where it provides the same resolution enhancement (Figure 5A-C; Figure S7; Movie S3).

**Figure 5.**
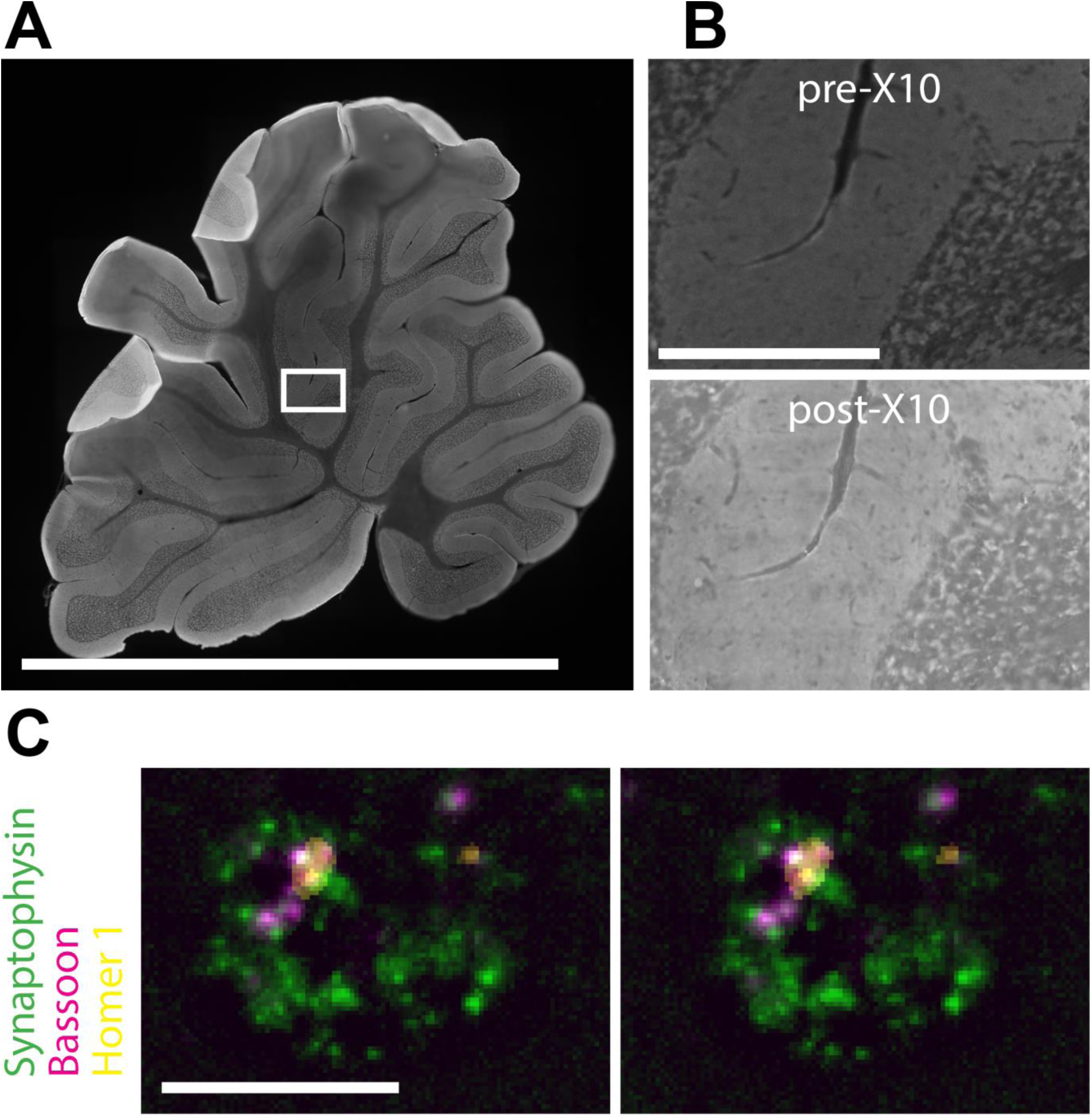
Multi-colour X10 imaging in brain slices. (A) Overview image of a rat cerebellum brain slice before expansion, showing Synaptophysin staining. Scale bar: 0.5 cm. Note that the image is stitched together from multiple imaging frames. (B) Magnification of the section framed in A), before expansion (top panel) and after expansion (bottom panel). Expansion factor: 9.6x. Scale bar: 250 μm. Note that both images are stitched together from multiple imaging frames. (C) A stereo view of a single synapse in the brain slice. Individual synaptic vesicles (identified by Synaptophysin, green), along with the active zone (identified by Bassoon, red) and the postsynaptic density (identified by Homer 1, yellow) are evident. Scale bar: 1 μm.

We therefore conclude that X10 microscopy provides the same resolution as reported in iterative expansion microscopy (iExM).^25^ The average resolution values are 25.2 nm for X10, and 25.8 nm for iExM. As indicated above, in iExM, the classical 4x gel is applied to the same sample multiple times in sequence, to achieve a multiplication of the expansion factors of the individual gels. This approach yields an expansion factor of up to ~16-20x, in two expansion steps. However, iExM is much more variable in its expansion factor than X10, as the multiplication of two 4x gels also results in a multiplication of the variability in their individual expansion factors, which usually spans from 3.5x to 4.5x.^7-10,25^ At the same time, the iterative protocol of iExM requires additional time and effort to break the first gel and prepare the second gel, and is not compatible (at the moment) with the use of conventional off-the-shelf antibodies, but requires custom-made DNA-oligo coupled antibodies.^25^ This makes iExM much more complex, more time-consuming, and less precise than X10.

That said, it may be possible in the future to combine the X10 gel with the iExM principle to achieve lateral expansion factors of up to 100x, which would theoretically offer a resolution of up to 3 nm, and which should be useful, if probes smaller than antibodies are employed. As the antibodies place the fluorophores at a substantial distance from the epitopes, the additional image detail obtained with expansion factors >10-12x comes at a disadvantage: the size of the measured structures no longer fits with the size of the actual objects measured, as shown by our simulations of microtubules (Figure S5), and by recent microtubule imaging results with iExM.^25^ The microtubule lumen can be resolved at expansion factors beyond ~15x using antibodies, but the apparent wall-to-wall distance of microtubules exceeds the expected microtubule diameter by 20-25 nm, averaging ~50-55 nm both in our simulations and in published iExM results.^25^ It is probably this factor that limits the effective resolution of iExM to ~25 nm, despite the larger expansion factor. X10, whose resolution is at the size of the probes, suffers less from this problem. Assuming that the Pmp70 epitopes surpass the peroxisome membrane by ~5 nm,^30^ and that the protein and antibody orientations are randomized after the permeabilization of the 6.8 nm-thick peroxisome membrane, the displacement induced by the antibodies is ~6 nm (Figure S4).

X10 thus provides a toolset for a cheap (less than 2$ for all reagents used in one experiment, except antibodies) and easy to use (no special machinery or custom antibodies required), multicolour super resolution imaging, which can be performed on widely available epifluorescence setups. Combining X10 with physics-based super-resolution, and especially with a coordinate-targeted approach such as STED,^31^ which can be applied rapidly and efficiently to large imaging volumes, would provide an ultimate resolution equal to the size of the fluorophores. This should enable the investigation of molecular assemblies or molecule orientation more efficiently than virtually any other current tools, especially if probes smaller than antibodies are employed.^13,14,21^

## ACKNOWLEDGMENT

We thank Martin Helm and Felipe Opazo for help with the STED imaging. S.T. was supported by an Excellence Stipend of the Göttingen Graduate School for Neurosciences, Biophysics, and Molecular Biosciences (GGNB). This work was supported by grants to S.O.R. from the European Research Council (ERC-2013-CoG NeuroMolAnatomy) and from the Deutsche Forschungsgemeinschaft (DFG): SFB 889/A5, 1967/7-1, and SFB1190/P09.

## ABBREVIATIONS

DMAA: *N,N*-dimethyl-acrylamide
FWHM: full width at half maximum
GSDIM: ground state depletion microscopy followed by individual molecule return
iExM: iterative expansion microscopy
KPS: potassium persulfate
PALM: photo-activated localization microscopy
STED: stimulated emission depletion
STORM: stochastic optical reconstruction microscopy
TEMED: *N,N,N’,N’*-tetramethyl-ethylene-diamine
X10: expansion microscopy with a 10-fold expansion factor.

## AUTHOR INFORMATION

The manuscript was written through contributions of all authors. All authors have given approval to the final version of the manuscript. S.T. and D.C. performed the stainings. M.M. cultured COS7 cells, produced the labelled scFv for Tubulin stainings, and performed the STED imaging. H.W. performed the STORM imaging. S.T. performed all other experiments and the expansion microscopy imaging. S.T., S.K., and S.O.R. conceived the project. S.T and S.O.R. performed the data evaluation and wrote the manuscript.

## REFERENCES

1. Hell, S.W.; Wichmann, J.. Breaking the diffraction resolution limit by stimulated emission: stimulated-emission-depletion fluorescence microscopy. Opt. Lett. 1994, 19, 780–782.

2. Willig, K.I.; Rizzoli, S.O.; Westphal, V.; Jahn R; Hell, S.W. STED microscopy reveals that synaptotagmin remains clustered after synaptic vesicle exocytosis. Nature 2006, 440, 935–939.

3. Betzig, E.; Patterson, G.H.; Sougrat, R.; Lindwasser, O.W.; Olenych, S.; Bonifacino, J.S.; Davidson, M.W.; Lippincott-Schwartz, J.; Hess, H.F. Imaging intracellular fluorescent proteins at nanometer resolution. Science 2006, 313, 1642–1645.

4. Rust, M.J.; Bates, M.; Zhuang, X. Sub-diffraction-limit imaging by stochastic optical reconstruction microscopy (STORM). Nat. Methods 2006, 3, 793–795.

5. van de Linde; S.; Löschberger, A.; Klein, T.; Heidbreder, M.; Wolter, S.; Heilemann, M.; Sauer, M. Direct stochastic optical reconstruction microscopy with standard fluorescent probes. Nat. Protoc. 2011, 6, 991–1009.

6. Testa, I.; Wurm, C.A.; Medda, R.; Rothermel, E.; von Middendorf, C.; Fölling, J.; Jakobs, S.; Schönle, A.; Hell, S.W.; Eggeling, C. Multicolor fluorescence nanoscopy in fixed and living cells by exciting conventional fluorophores with a single wavelength. Biophys. J. 2010, 99, 2686–2694.

7. Chen, F.; Tillberg, P.W.; Boyden, E.S. Optical imaging. Expansion microscopy. Science 2015, 347, 543–548.

8. Chozinski, T.J.; Halpern, A.R.; Okawa, H.; Kim, H.J.; Tremel, G.J.; Wong, R.O.; Vaughan, JC. Expansion microscopy with conventional antibodies and fluorescent proteins. Nat. Methods 2016, 13, 485–488.

9. Chen, F.; Wassie, A.T.; Cote, A.J.; Sinha, A.; Alon, S.; Asano, S.; Daugharthy, E.R.; Chang, J.B.; Marblestone, A.; Church, G.M.; Raj, A.; Boyden, E.S. Nanoscale imaging of RNA with expansion microscopy. Nat. Methods 2016, 13, 679–684.

10. Tillberg, P.W.; Chen, F.; Piatkevich, K.D.; Zhao, Y.; Yu, C.C.; English, B.P.; Gao, L.; Martorell, A.; Suk, H.J.; Yoshida, F.; DeGennaro, E.M.; Roossien, D.H.; Gong, G.; Seneviratne, U.; Tannenbaum, S.R.; Desimone, R.; Cai, D.; Boyden, E.S. Protein-retention expansion microscopy of cells and tissues labeled using standard fluorescent proteins and antibodies. Nat. Biotechnol. 2016, 34, 987–992.

11. Cipriano, B.H.; Banik, S.J.; Sharma, R.; Rumore, D.; Hwang, W.; Briber, R.M.; Raghavan, S.R. Superabsorbent hydrogels that are robust and highly stretchable. Macromolecules 2014, 47, 4445–4452.

12. Zhao, Y.; Bucur, O.; Irshad, H.; Chen, F.; Weins, A.; Stancu, A.L.; Oh, E.Y.; DiStasio, M.; Torous, V.; Glass, B.; Stillman, I.E.; Schnitt, S.J.; Beck, A.H.; Boyden, E.S. Nanoscale imaging of clinical specimens using pathology-optimized expansion microscopy. Nat. Biotechnol. 2016, 35, 757–764.

13. Ries, J.; Kaplan, C.; Platonova, E.; Eghlidi, H.; Ewers, H. A simple, versatile method for GFP-based super-resolution microscopy via nanobodies. Nat. Methods 2012, 9, 582–584.

14. Fornasiero, E.F.; Opazo, F. Super-resolution imaging for cell biologists: concepts, applications, current challenges and developments. Bioessays 2015, 37, 436–451.

15. Wildanger, D.; Rittweger, E.; Kastrup, L.; Hell, S.W. STED microscopy with a supercontinuum laser source. Opt. Express 2008, 16, 358–362.

16. Wildanger, D.; Bϋckers, J.; Westphal, V.; Hell, S.W.; Kastrup, L. A STED microscope aligned by design. Opt. Express 2009, 17, 209–213.

17. Wegel, E.; Göhler, A.; Lagerholm, B.C.; Wainman, A.; Uphoff, S.; Kaufmann, R.; Dobbie, I.M. Imaging cellular structures in super-resolution with SIM, STED and Localisation Microscopy: A practical comparison. Sci. Rep. 2016, 6, 27290.

18. Niehörster, T.; Löschberger, A.; Gregor, I.; Krämer, B.; Rahn, H.J.; Patting, M.; Koberling, F.; Enderlein, J.; Sauer, M. Multi-target spectrally resolved fluorescence lifetime imaging microscopy. Nat. Methods 2016, 13, 257–262.

19. Bálint, Š.; Verdeny-Vilanova, I.; Sandoval-Álvarez, Á.; Lakadamyali, M. Correlative live-cell and superresolution microscopy reveals cargo transport dynamics at microtubule intersections. Proc. Natl. Acad. Sci. U. S. A. 2013, 110, 3375–3380.

20. Bates, M.; Huang, B.; Dempsey, G.T.; Zhuang, X. Multicolor super-resolution imaging with photo-switchable fluorescent probes. Science 2007, 317, 1749–1752.

21. Mikhaylova, M.; Cloin, B.M.; Finan, K.; van den Berg, R.; Teeuw, J.; Kijanka, M.M.; Sokolowski, M.; Katrukha, E.A.; Maidorn, M.; Opazo, F.; Moutel, S.; Vantard, M.; Perez, F.; van Bergen en Henegouwen, P.M.; Hoogenraad, C.C.; Ewers, H.; Kapitein, L.C. Resolving bundled microtubules using anti-tubulin nanobodies. Nat. Commun. 2015, 6, 7933.

22. Huang, B.; Jones, S.A.; Brandenburg, B.; Zhuang, X. Whole-cell 3D STORM reveals interactions between cellular structures with nanometer-scale resolution. Nat. Methods 2008, 5, 1047–1052.

23. Dani, A.; Huang, B.; Bergan, J.; Dulac, C.; Zhuang, X. Superresolution imaging of chemical synapses in the brain. Neuron 2010, 68, 843–856.

24. Schmidt, R.; Wurm, C.A.; Jakobs, S.; Engelhardt, J.; Egner, A.; Hell, S.W. Spherical nanosized focal spot unravels the interior of cells. Nat. Methods 2008, 5, 539–544.

25. Chang, J.B.; Chen, F.; Yoon, Y.G.; Jung, E.E.; Babcock, H.; Kang, J.S.; Asano, S.; Suk, H.J.; Pak, N.; Tillberg, P.W.; Wassie, A.T.; Cai, D.; Boyden, E.S. Iterative expansion microscopy. Nat. Methods 2017, 14, 593–599.

26. Balzarotti, F.; Eilers, Y.; Gwosch, K.C.; Gynnå, A.H.; Westphal, V.; Stefani, F.D.; Elf, J.; Hell, S.W. Nanometer resolution imaging and tracking of fluorescent molecules with minimal photon fluxes. Science 2017, 355, 606–612.

27. Gao, R.; Asano, S.M.; Boyden, E.S. Q&A: Expansion microscopy. BMC Biol. 2017, 15, 1–9.

28. Glebov, O.O.; Cox, S.; Humphreys, L.; Burrone, J. Neuronal activity controls transsynaptic geometry. Sci. Rep. 2016, 6, 22703.

29. Tang, A.H.; Chen, H.; Li, T.P.; Metzbower, S.R.; MacGillavry, H.D.; Blanpied, T.A. A trans-synaptic nanocolumn aligns neurotransmitter release to receptors. Nature 2016, 536, 210–214.

30. Baker, A.; Carrier, D.J.; Schaedler, T.; Waterham, H.R.; van Roermund, C.W.; Theodoulou, F.L. Peroxisomal ABC transporters: functions and mechanism. Biochem. Soc. Trans. 2015, 43, 959–965.

31. Hell, S.W. Far-field optical nanoscopy. Science 2007, 316, 1153–1158.

32. Strachota, B.; Matějka, L.; Zhigunov, A.; Konefal, R.; Spěváček, J.; Dybal, J.; Puffr, R. Poly(N-isopropylacrylamide)-clay based hydrogels controlled by the initiating conditions: evolution of structure and gel formation. Soft Matter, 2015, 11, 9291–9306

33. Carlsson, L.; Rose, S.; Hourdet, D.; Marcellan, A. Nano-hybrid self-crosslinked PDMA/silica. Soft Matter, 2010, 6, 3619–3631

